# Sample preparation and data collection for Serial Block Face Scanning Electron Microscopy of Mammalian Cell Monolayers

**DOI:** 10.1101/2023.09.26.559595

**Authors:** Noelle V. Antao, Joseph Sall, Christopher Petzold, Damian C. Ekiert, Gira Bhabha, Feng-Xia Liang

## Abstract

Volume electron microscopy encompasses a set of electron microscopy techniques that can be used to examine the ultrastructure of biological tissues and cells in three dimensions. Two block face techniques, focussed ion beam scanning electron microscopy (FIB-SEM) and serial block face scanning electron microscopy (SBF-SEM) have often been used to study biological tissue samples. More recently, these techniques have been adapted to *in vitro* tissue culture samples. Here we describe detailed protocols for two sample embedding methods for *in vitro* tissue culture cells intended to be studied using SBF-SEM. The first protocol focuses on cell pellet embedding and the second on *en face* embedding. *En face* embedding can be combined with light microscopy, and this CLEM workflow can be used to identify specific biological events in a light microscope, which can then be imaged using SBF-SEM. We systematically outline the steps necessary to fix, stain, embed and image adherent tissue culture cell monolayers by SBF-SEM. In addition to sample preparation, we discuss optimization of parameters for data collection. We highlight the challenges and key steps of sample preparation, and the consideration of imaging variables that will facilitate the acquisition of high quality datasets. Users experienced with electron microscopy sample preparation methodology will be able to complete this protocol in 10-11 days from initial seeding of cells in tissue culture to image acquisition.

## INTRODUCTION

Volume electron microscopy (vEM) can be used to generate 3D reconstructions of biological samples ranging from tissues to individual cells at resolutions higher than light microscopy^1^. In vEM, a series of 2-dimensional images are collected, which can subsequently be processed, annotated and reconstructed in 3 dimensions. Typically, vEM techniques provide spatial resolutions at which the interior structures of organelles can be deciphered^2^, for example, the cristae of mitochondria. Two block-face scanning electron microscopy (SEM)-based techniques are predominantly used to image biological samples in 3D: focused ion beam scanning electron microscopy (FIB-SEM)^3^ and serial block-face scanning electron microscopy (SBF-SEM)^4^. In both methods, 3D volumes are acquired by serial sectioning of samples, using either a focused ion beam (FIB-SEM) or an ultramicrotome (SBF-SEM) to remove a thin layer of the sample automatically, followed by imaging of the newly exposed sample block surface by SEM. Although both techniques have a similar spatial resolution in the *x*-*y* dimension (3-5 nm), FIB-SEM has greater resolution in the *z* dimension for achieving isotropic voxels down to 3-5 nm voxel size for biological samples, due to the ability to cut thinner sections using a focused ion beam^5^. In contrast, resolution in the *z* direction is somewhat lower in SBF-SEM, as the thinnest section that can be reliably cut using an ultramicrotome is ∼25 nm in thickness^6^. While the *z* resolution is higher using FIB-SEM, the milling process is slower, which can, in practice, limit the total area and volume that can be imaged relative to SBF-SEM. The ultramicrotome/diamond knife used for sectioning in SBF-SEM makes the cutting of sections about 4 times faster than ion beam milling, enabling efficient imaging of samples with larger volumes^2,7–9^.

The SBF-SEM instrument is comprised of an SEM, an ultramicrotome mounted within the SEM chamber, and the necessary hardware and software to control image acquisition^4^. The first imaging setup resembling modern SBF-SEM was invented in 1981 by SB Leighton, in which a miniature microtome mounted within the SEM chamber was used to cut and image successive sections from a resin embedded squid fin nerve tissue sample^10^. Limitations of this first setup included: 1) The need to remove the sample from the chamber in between sections, in order to carbon coat it for optimal conductivity, and 2) The limited capabilities for storage and analysis of digital image files, given existing technology at the time. The technique was revisited and improved upon by Denk and Horstmann^4^, who used a variable pressure system, which allowed imaging and analysis of non-conductive samples, overcoming the need for carbon coating between sections. Since then, SBF-SEM has been used to examine biological structures at different length scales from micrometers (axons) to nanometers (individual chromosomes and mitochondrial cristae) ^7–9^.

Historically, SBF-SEM has been extensively used to study biological tissue samples such as brain tissue for neuronal circuit reconstruction^11,12^. More recently this methodology has been adapted to study the ultrastructure of other biological tissue samples e.g. bone tissue, developing unicellular organisms and cell monolayers. The varying characteristics of different biological samples requires that sample preparation and image acquisition be tailored to the biological sample being studied. Here, we systematically address the specific challenges associated with sample preparation and imaging of *in vitro* cultured cells by SBF-SEM. We outline all the steps necessary to stain, embed and image *in vitro* cultured monolayer cells for SBF-SEM analysis. We also outline the steps necessary for performing correlative light and electron microscopy (CLEM) in order to target specific or rare biological events^13,14–15,16^ prior to SBF-SEM data collection. The steps detailed in the protocol are optimized for the analysis of adherent mammalian cell lines using the Gatan OnPoint BSE detector in a Zeiss Gemini300 VP FESEM equipped with a Gatan 3View automatic ultramicrotome with a focus charge compensator (FCC) installed to reduce sample charge accumulation during imaging acquisition. The basic methodologies detailed in this protocol can also be adapted to other cell types and vEM methodologies.

Our target audience for this protocol are newcomers to SBF-SEM, who may benefit from an introduction to optimizing sample preparation and imaging conditions for SBF-SEM data collection of *in vitro* cultured cell samples as well as more experienced microscopists planning to image cellular samples for the first time.

### Overview of SBF-SEM

The goal for vEM imaging techniques such as SBF-SEM is to obtain quantitative volumetric information of biological samples with the best image quality and highest resolution possible. It is important that the data be of highest quality to ensure the reliability and robustness of 3D reconstructions and quantification, and to facilitate automation of data analysis, such as image segmentation. In SBF-SEM imaging, image quality is determined by: 1) image contrast (signal-to-noise), which can be visualized as differences in electron densities; 2) image resolution, which is defined by pixel size and can be as low as 2 nm;^17^ and 3) cutting stability, which refers to the uniformity with which each slice is being cut. The important steps that require optimization to ensure reliable generation of volumetric image data by SBF-SEM can broadly be divided into 1) sample preparation and 2) image acquisition.

#### Sample preparation

High quality SBF-SEM data acquisition relies on robust sample preparation. SBF-SEM samples must be prepared such that they: 1) preserve the ultrastructure of the biological tissue, 2) have the structural rigidity to withstand the high vacuum and ultramicrotome slicing within the SEM chamber, 3) have sufficient contrast during imaging and 4) optimize sample conductivity to reduce charging artifacts during imaging. The first step in the sample preparation pipeline is tissue fixation and staining, which is most commonly a variation of the Deernick method^18,19–20^. In this method, the biological tissue sample is treated with chemical fixatives; for example, a mixture of aldehydes, and then with multiple heavy metal stains simultaneously (*en bloc)* to enhance the contrast of the sample as well as improve sample conductivity. These heavy metal salts include: osmium tetroxide, which preferentially interacts with unsaturated lipids; thiocarbohydrazide, which binds to osmium in the tissue and acts as a bridging agent to which more osmium can bind, thereby enhancing the contrast of lipid components in the cell; uranyl acetate, which stains nucleic acids and proteins an provides general contrast to biological samples; lead aspartate, which interacts with membranes and proteins. Following fixation and staining, samples are dehydrated by incubation in a series of graded organic solvent solutions, such as ethanol, and then embedded in an epoxy resin^21,22^. The samples are next trimmed to a shape (truncated pyramid, cube or cuboid) and size that will be amenable to cutting by the ultramicrotome mounted in the SEM chamber. Depending on the biological sample being analyzed, each of the following steps in sample preparation require optimization at the level of: 1) the combination and concentration of heavy metals, 2) the incubation time of each stain to ensure even stain penetration throughout the sample, 3) the type of embedding resin and 4) the dimensions of the final sample block that is loaded into the SEM chamber. Special fixation and sample preparation methods such as high pressure freezing and freeze substitution should be considered to analyze fast cellular events. In this protocol, we discuss the two most popular methods for preparing *in vitro* cultured cell samples: 1) cell pellets (Figure 1A) and 2) *en face* monolayer cells (Figure 1B).

**Figure 1:**
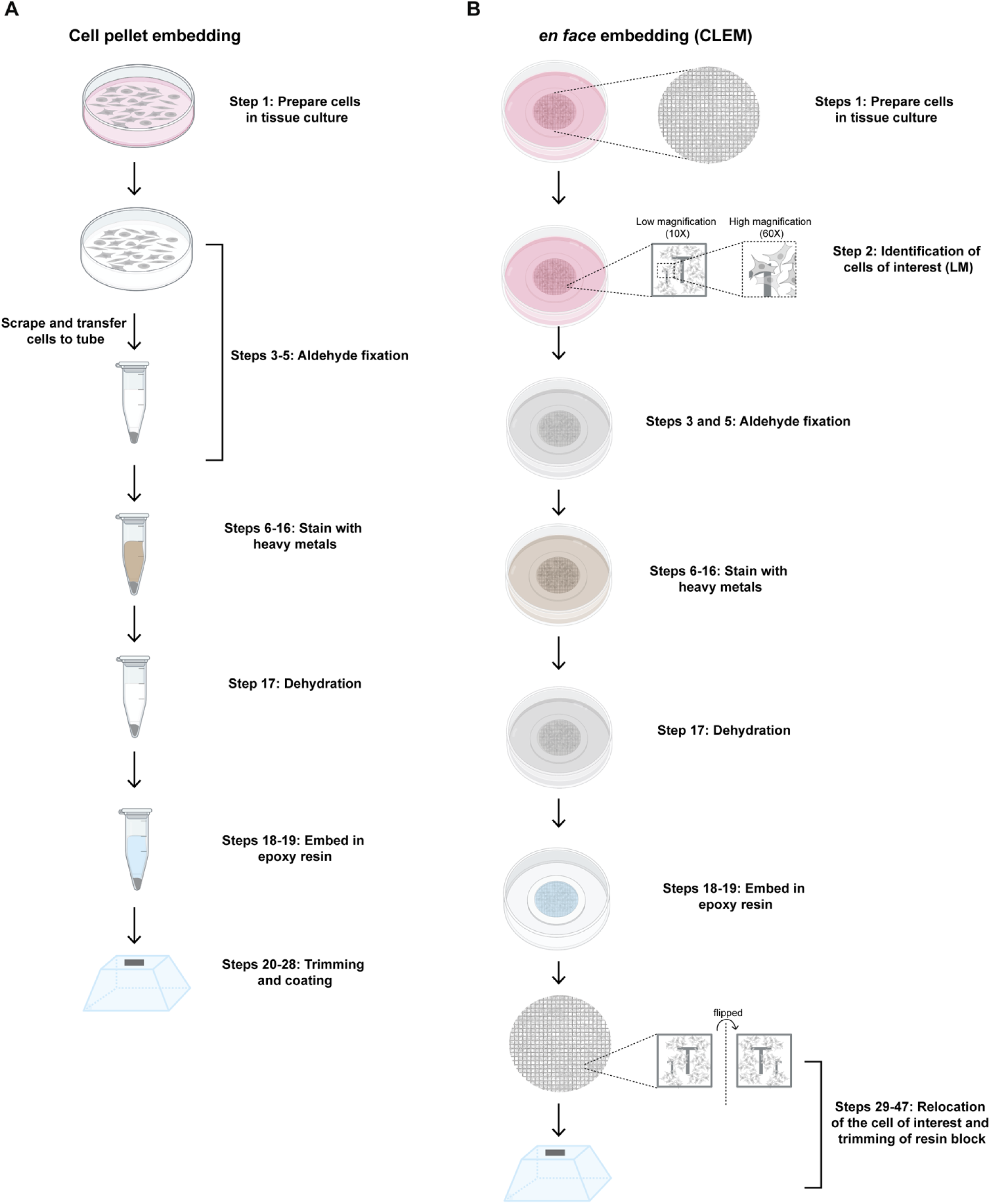
Key steps in the workflow for SBF-SEM of adherent tissue culture cells. (A) Cell pellet workflow and (B) *en face* with CLEM workflow. Created in part with Biorender. com

#### Image acquisition

The imaging cycle begins with an electron beam scanning across the blockface to generate images from high energy backscattered electrons (BSE’s) emitted from the sample. Once the entire region of interest is imaged, the sample block is raised by a specified distance, corresponding to slice thickness, in the z-direction. The ultramicrotome then removes a thin section from the surface and the newly exposed blockface is imaged. This sequence of imaging steps is repeated to generate a series of images that are aligned in *x-y* and registered in the z-direction (Figure 2A). A significant challenge during image acquisition is sample charging that results from an accumulation of low energy secondary electrons within the sample blockface. This build-up of charge damages the sample blockface, distorts acquired images, and impacts image contrast. Several imaging parameters influence image contrast and cut stability during acquisition, including accelerating voltage, aperture size, dwell time, pixel size and focal charge compensation (Figure 2B). Varying each of these parameters affects different aspects of the resulting images; for example, increasing accelerating voltage increases signal to noise, but higher accelerating voltages can also compromise the block face, resulting in charging artifacts and distortion of the acquired images (Figure 2C). Charging artifacts can be resolved by altering parameters like beam current, dwell time and pixel size during image acquisition (Figure 2C). Additionally, a focus charge compensator (FCC) is installed very close to the sample block face, so that ionized nitrogen gas molecules can neutralize the charge build up on the sample surface, generated by secondary electrons^23,24–25^ (Figure 2C). While increasing the nitrogen gas pressure and/or decreasing dwell time will compensate for charging and reduce charging artifacts, it does come at a cost to signal-to-noise. Therefore, it is important to tailor imaging parameters to identify image acquisition conditions that maximize the ability to acquire data at the highest resolution without compromising signal-to-noise or cut stability.

**Figure 2:**
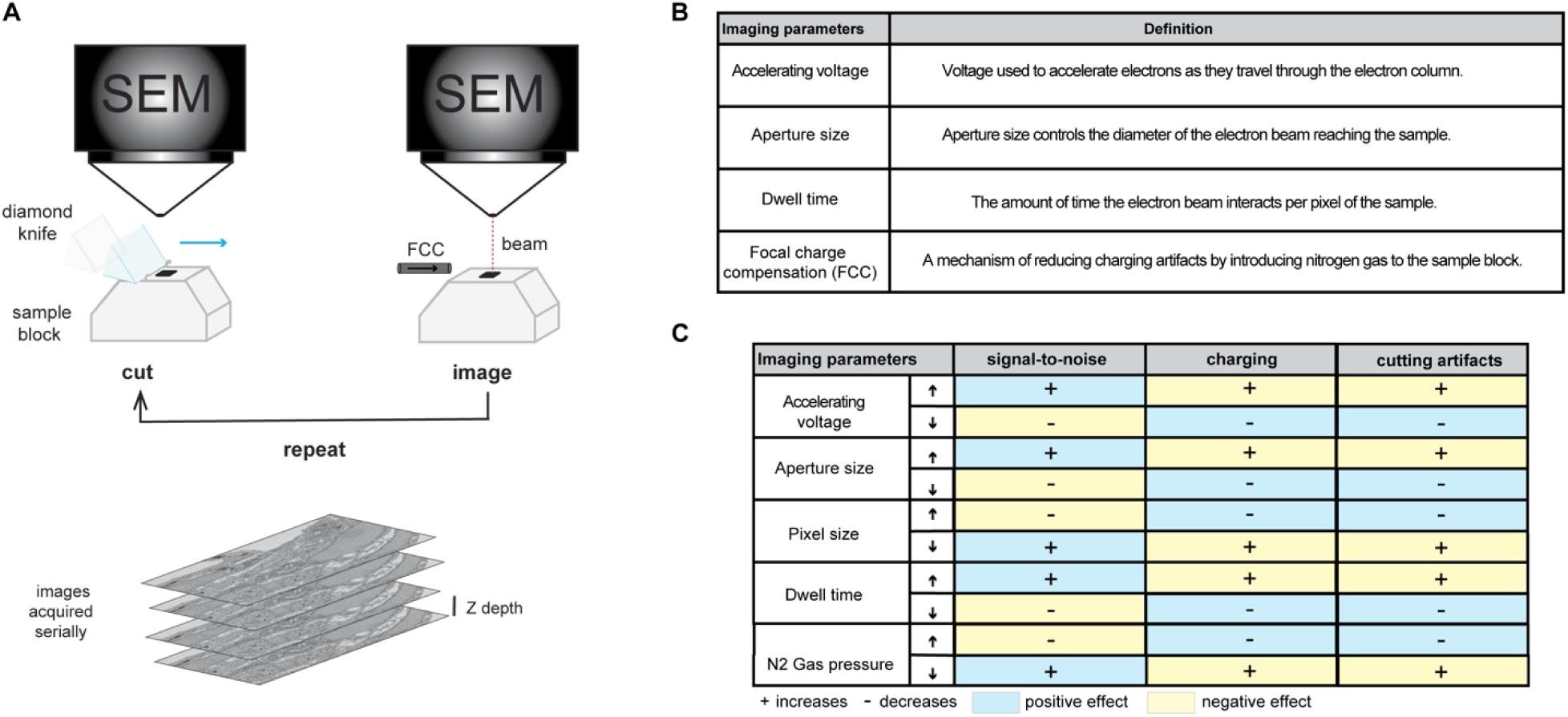
SBF-SEM imaging concept and key imaging parameters. (A) Schematic of the SBF-SEM sample and imaging setup. (B) Table of the key imaging parameters and their definitions. (C) Table describing how increasing (up arrow) or decreasing (down arrow) key parameters impacts properties of the resulting image, including signal-to-noise, charging and cutting artifacts during imaging.

### Embedding and imaging tissue culture monolayers using SBF-SEM

Life science research relies heavily on mammalian cell lines cultured as monolayers *in vitro*. However, it can be challenging to image cell monolayers by SBF-SEM, in large part due to their limited depth, and the presence of large areas of non-conductive resin between cells, an issue not present when preparing the majority of tissue samples. The absence of many conductive paths in these samples can result in the buildup of negative charge at the sample surface leading to cutting artifacts and low image quality. Therefore, SBF-SEM imaging of cell monolayers requires modifications of standard sample preparation protocols not only in sample staining and embedding, but also on sample mounting methods, as well as an adjustment of the different imaging parameters to ensure robust imaging and data quality. Previously, SBF-SEM has been used to examine the organization of organelles as well as to characterize the life cycle of pathogens in tissue culture monolayers^26–31,14^. Depending on the biological question being addressed, two sample preparation techniques are particularly useful 1) embedding as cell pellets or 2) *en face* embedding of cell monolayers. In the embedded cell pellet method, adherent cells are first fixed and detached from the cell culture dish, and pelleted, prior to staining and embedding (Figure 1A). In the *en face* embedding method, cells are fixed, stained and embedded directly in the cell culture dish thereby preserving the orientation of the cell (Figure 1B). The cell pellet method is ideal when the underlying orientation of the cell is not important and the biological event being studied is observed frequently (ideally in >50% of cells). Because cells in the pellet are tightly packed together, many cells can be imaged simultaneously. The *en face* method is particularly useful to study rare biological events by correlative microscopy, as it allows users to combine the information from two different microscopy modalities - light microscopy (LM) to identify the rare event, and electron microscopy (SBF-SEM) to image it at high resolution. However, with a limited field of view, the imaging area is restricted to just one or a few cells at a time, and therefore the throughput of *en face* embedding and imaging is low. For any given sample, the choice of cell pellet embedding versus *en face* embedding will depend on the underlying biology being examined. A list of advantages and disadvantages for each sample preparation method are described in Table 2.

**Table 1:** Table of technical specifications for a Zeiss Gemini300 SEM with a 3View2XP.

| Parameter | GeminiSEM 300 with a 3View2XP |
| --- | --- |
| Resolution | 0.8 nm at 15kV; 1.4 nm at 1kV |
| Accelerating Voltage | <1kV - 5kV |
| Probe Current | 3pA - 20 nA |
| Charge Neutralization | 5 - 40 Pa is typical at 2kV |
| Cut Speed | Recommended speed: 0.5 - 1 mm/s |
| Cut Thickness | 25 - 50 nm to analyze cell structure |
| Pixel Dwell Time | 1 -20 μs |
| Image Scanning | Single pass or multiple frames |
| Imaging modes | Single frame per cut; Multiple fields of view and magnifications per cut; stage montage for large fields of view |

**Table 2:** Table of advantages and disadvantages of the cell pellet versus *en face* embedding of adherent tissue culture cells.

| Method | Pro | Con |
| --- | --- | --- |
| Cell pellet | Higher throughput (multiple cells or events per sample block) | Complete cells are difficult to capture |
|  | Amenable to adherent and suspension cell cultures | Requires that the biological event being studied occurs frequently (>50% of cells) |
|  | Increased sample depth for imaging |  |
|  | Better sample conductivity owing to crowded or densely packed cells pellets |  |
| *en face* with CLEM | Identify and capture rare biological events | Lower throughput (1 cell per sample block) |
|  | Complete cell can be captured | Poorer conductivity owing to limited sample depth |

### Advantages and limitations of SBF-SEM

The advantage of SBF-SEM lies in the ability of this technique to acquire quantitative ultrastructural information on biological samples in an automated manner over large imaging areas and volumes^32–34^. Image acquisition is fully automated, and serially acquired images are well-aligned, with small amounts of image translation in the x and y that can be corrected post acquisition. Moreover, correlative light and SBF-SEM (CLEM) allows tracking of specific biological events or subcellular location by either widefield or fluorescence microscopy and correlates it to the underlying ultrastructure in the area of interest. Limitations to this technique are similar to all EM ultrastructural analysis, 1) sample preparation takes several days and should be tailored to the biological sample being studied, 2) the use of chemical fixatives and heavy metal staining during sample preparation may alter the properties of the underlying ultrastructure. Alternatively, high pressure freezing and freeze substitution can be considered in order to capture fast cellular events and have better structural preservation, and 3) z-resolution is limited by the ultramicrotome knife (∼25 nm) ^29^. Depending on the biological question, other vEM techniques might also be useful to consider, such as FIB-SEM^3^ which has greater Z-resolution, array tomography^35^, which can be combined with protein localization, and TEM-based vEM such as serial section TEM^36^ (ssTEM) and serial section electron tomography (ssET)^37^ with higher *x-y* resolution.

### Level of expertise needed to implement this protocol

Users should be experienced with standard tissue culture practices, EM sample preparation equipment/accessories, and have access to SBF-SEM instrumentation, as well as staff with the technical expertise to operate the instrument. For further image analysis beyond this protocol, users should have available software for image alignment and 3D rendering, such as Fiji^38^ and Dragonfly software^39^ or its equivalent.

## EXPERIMENTAL DESIGN

Below we outline key steps to consider when planning an SBF-SEM experiment with adherent cells.

### Sample preparation

#### Mammalian cell culture for cell pellets

Adherent cells can be grown on regular tissue culture dishes and suspension cells can be grown in flasks prior to sample staining and embedding.

#### Mammalian cell culture for *en face* samples for CLEM

Adherent mammalian cells must be grown on a gridded coverslip, for example, a 35 mm dish with a gridded coverslip (MatTek Cat no. P35G-1.5-14-CGRD). Cell seeding densities for CLEM should be optimized to prevent the growth of dense cell monolayers, which will obscure the underlying grid numbers during light microscopy (step 3). It is important that the grid numbers are visible for identifying the relevant region of interest during SBF-SEM imaging.

#### Selecting a Fixation Method

When selecting a sample fixation method, it is important to consider the type of biological event being evaluated as well as sample size. In general, a chemical fixation that combines different ratios of aldehydes (2-4% PFA and 1-5% glutaraldehyde) should meet the majority of fixation requirements. However, cryo fixation with high pressure freezing and freeze substitution is recommended when trying to capture rapid biological events, or when it is especially important to preserve specific biological structures.

#### *En bloc* staining

Depending on each project, a different combination of heavy metal stains may be optimal. For example, osmium tetroxide is an essential heavy metal stain with biological materials as it stains unsaturated lipids on the plasma membrane. Additionally, combining osmium tetroxide staining with thiocarbohydrazide helps increase not only the contrast, but also the conductivity of most biological samples.

However, osmium tetroxide is not always suitable particularly when trying to preserve biological structures like the cytoskeleton. In such cases, using heavy metals like potassium ferri- or ferrocyanide might be preferred. Alternatively, the addition of heavy metals like uranyl acetate, lead aspartate and tannic acid can also be considered when trying to improve the contrast of biological samples.

#### Dehydration and embedding

A graded series of organic solvents such as ethanol or acetone are used to replace the water in biological samples through dehydration during sample preparation. Acetone is a stronger organic solvent and mixes well with epoxy resin, however, it is not an ideal solvent for *en face* embedding and sample processing as it will dissolve the plastic dish. There are several epoxy resins that can be chosen for SBF-SEM sample embedding, such as Durcupan, hard EMbed 812 and Spurr. We do not recommend EMbed 812 resin for *en face* samples as it might dissolve the plastic dish thereby compromising sample recovery.

#### Sample trimming and mounting

Either manual or machine trimming can be used when removing excess resin. For cell pellet samples, the removal of excess resin is important to expose both the top surface of the sample (blockface that will be imaged) as well as the bottom surface, if possible, which is glued to the 3View pin stub. The use of a silver conductive epoxy glue during this process allows for improved charge dissipation during SEM imaging by grounding the sample to the 3View pin. For *en face* embedded CLEM samples, removing as much excess resin from the non-sample side that is glued to the 3View pin is critical for successful SBF-SEM imaging as it grounds the sample and helps with charge dissipation (Figure 3). Specimens should be inspected for the quality of sample preservation and the presence of any preparation artifacts such as sample deformation or changes in organelle ultrastructure via TEM. For *en face* samples, users will likely need to sacrifice a cell or region of interest in order to verify the quality of sample staining and the preservation of organelle ultrastructure.

**Figure 3:**
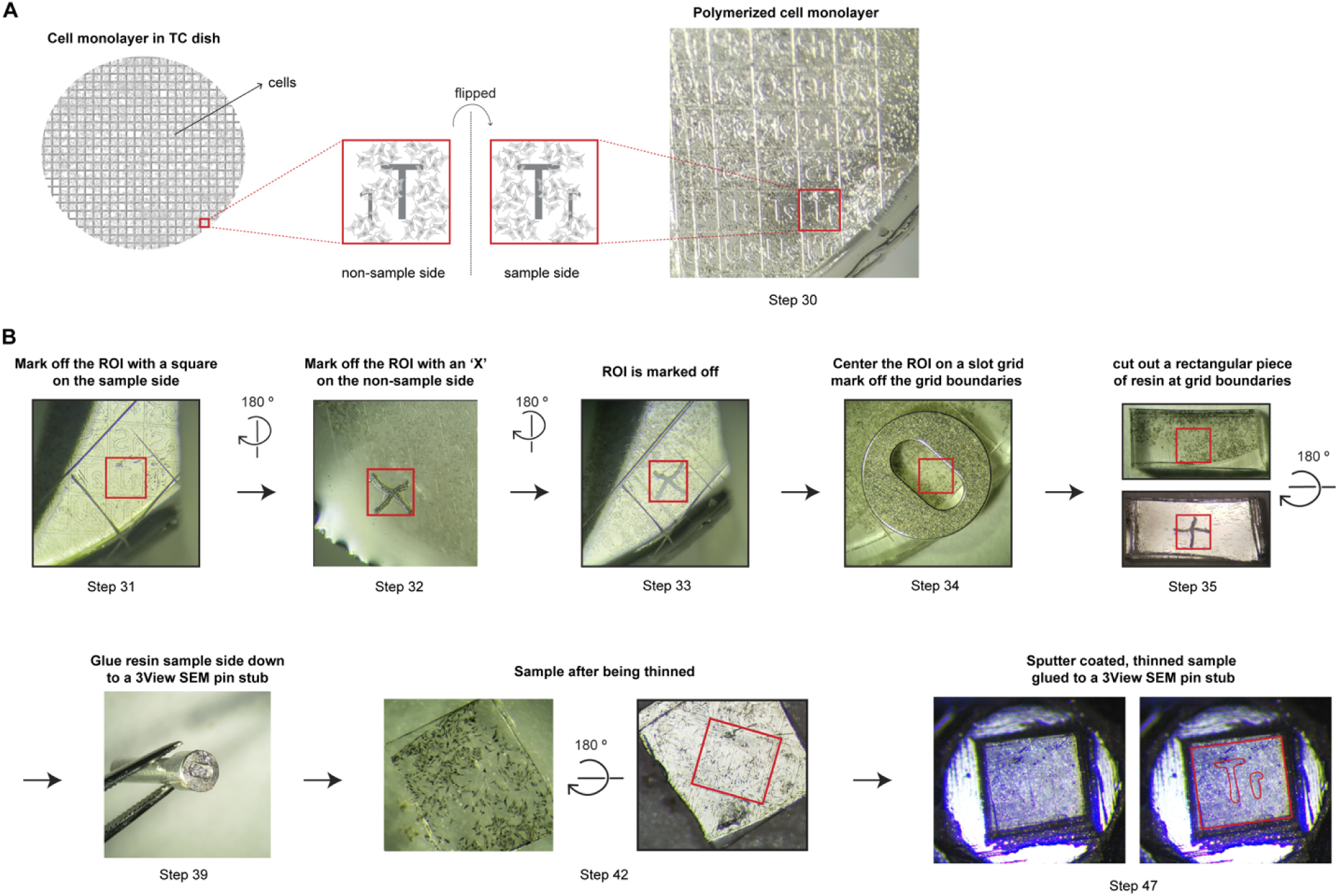
Key steps in trimming and mounting of *en face* embedded samples. (A) Image of an *en face* embedded sample with the ROI highlighted. (B) Pictures highlighting the steps in the workflow for trimming and mounting of the ROI highlighted in (A) onto a 3View pin stub. Steps noted in the figure correspond to steps of the protocol. Created in part with Biorender.com.

### SBF-SEM instrumentation

SBF-SEM uses an *in situ* ultramicrotome (diamond knife) mounted inside an SEM chamber to cut thin sections of the sample block (Figure 4). Following each cutting event, the sample block is raised by a specified distance (*z* slice thickness) and imaged. This cutting and imaging cycle repeats automatically until the desired volume is collected. Achieving the proper resolution for most biological events requires a scanning electron microscope equipped with a field emission gun, and backscattered and secondary electron detectors designed for imaging at low accelerating voltages (1 - 2 keV). SBF-SEM collections can run for hours, weeks or months, depending on the volume and resolution being collected for a given project. Given the duration of these imaging experiments, the microscope requires anti-vibration hardware, magnetic field cancellation systems and minimal ambient temperature fluctuations.

**Figure 4:**
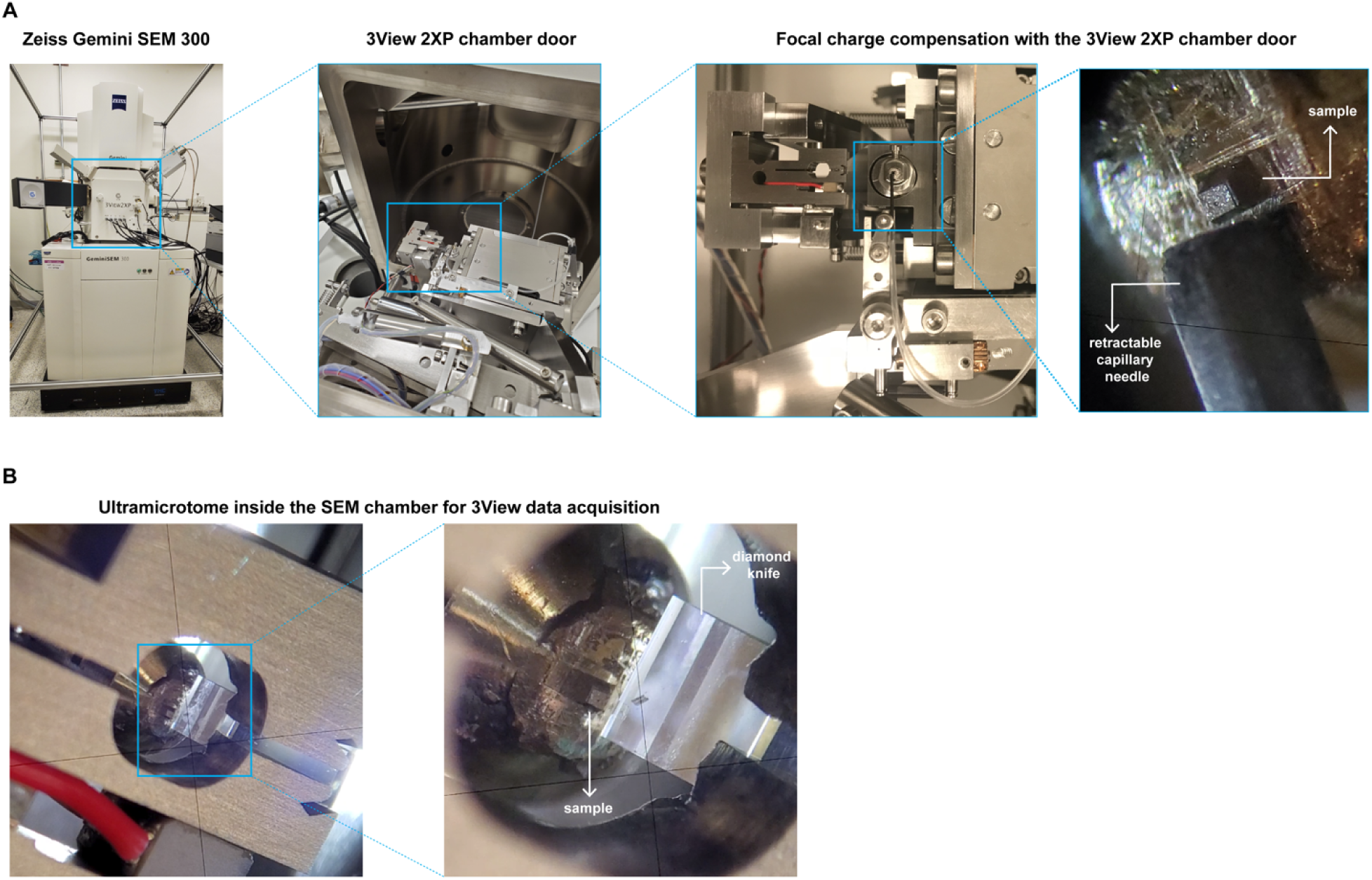
GeminiSEM 300 with a 3View2XP setup. (A) 3View2XP stage setup. Insets show the retractable nozzle for the Focal Charge Compensation system and the sample at higher magnification. (B) Picture of the retractable ultramicrotome system in the 3View2XP stage.

#### Preparation for SBF-SEM

After the sample is loaded into the SEM, the sample should be manually approached with the ultramicrotome, similar to TEM sectioning. For CLEM, the light microscopy images should be used as guidelines to select the area of interest for imaging.

#### Considerations for SBF-SEM Data Acquisition and Interpretation

Important considerations before SBF-SEM imaging include 1) image pixel size (resolution in *x-y*), 2) the *z* resolution (thickness of each slice) and 3) the volume (total number of slices) being collected. *x-y* dimensions can range from 20 µm to 80 µm, providing a maximum area of 80 µm x 80 µm. Multiple areas of interest can be collected sequentially for cell pellet samples. Depending on the biological questions being asked, a set of image acquisition parameters should be chosen including the accelerating voltage, probe current, pixel size, dwell time and scan speed (Figure 2B-C). Generally, a 5 - 6 nm pixel size is sufficient for resolving cell organelle ultrastructure. Monitor imaging periodically to adjust any focus drift that may occur during acquisition. Once data is collected, the serial images can be aligned and segmented using either open source or commercial software.

## MATERIALS

### Biological materials

In this protocol we use Vero cells (African green monkey epithelial) infected with *Encephalitozoon intestinalis*, a microsporidian parasite.

### Reagents

- Dulbecco’s Modified Eagle Medium, high glucose (DMEM:HG) (Thermo Fisher Scientific, Cat no. 11995065)
- Dulbecco’s phosphate-buffered saline (DPBS) (Thermo Fisher Scientific, Cat no. 14190144)
- Trypsin-EDTA (Thermo Fisher Scientific, Cat no. MT25053CI)
- Trypan Blue (Thermo Fisher Scientific T10282)
- Fetal Bovine Serum (VWR 89510-188)
- 100X Non essential amino acids (Fisher Scientific 11-140-050)
- Milli Q water
- 16% paraformaldehyde (Electron Microscopy Sciences, EMS, Cat no. 15700)
- 25% glutaraldehyde (Electron Microscopy Sciences, Cat no. 16019)
- 0.4 M sodium cacodylate buffer, pH 7.2 (Electron Microscopy Sciences, Cat no. 11654)
- 4% Osmium Tetroxide (Electron Microscopy Sciences, Cat no. 19150)
- Thiocarbohydrazide
- L-aspartic acid (Sigma-Aldrich)
- Lead nitrate (Ted Pella Inc., Cat no. 19321)
- Ethanol, 200 proof (Electron Microscopy Sciences, Cat no. 15056)
- Durcupan ACM Epoxy (Electron Microscopy Science, Cat no. 14040)
- EMbed 812 Embedding Kit (Electron Microscopy Science, Cat no. 14121)
- Silver conductive epoxy (Chemtronics CW2400 Epoxy)

### Equipment

- Biosafety cabinet
- CO_2_ incubator (Heracell CO_2_ Incubators, Thermo Scientific)
- Inverted light microscope to check confluence of cells (Carl Zeiss Microscopy LLC)
- Countess II Cell Counter (Thermo Fisher Scientific)
- Invitrogen Countess Cell Counting Chamber Slides (Thermo Fisher Scientific, Cat no. C10312)
- Ibitreat, 35mm tissue culture treated dish (Ibidi, Cat no. 81156)
- 35 mm tissue culture dish; No. 1.5 Gridded Coverslip; 14 mm Glass Diameter (MatTek Corporation, Cat no. P35G-1.5-14-CGRD)
- Cell scraper (VWR 10062-904)
- Wood Applicator Sticks (Solon, Cat no. FBHOO360)
- Embedding mold (PELCO, Cat no. 10535)
- Pipette and pipette tips
- Chemical hoods
- Razor (EMS Cat# 72000-WA)
- Formar Support Copper Slot Grid (Electron Microscopy Science, Cat no. FF2010)

### Specific equipment

- 3View SEM pin stub: 1.4mm flat, 2mm pin, 12mm height (EMS catalog# 75959-02)
- 3View SEM pin stub: 2.4mm flat, 2mm Pin, 12.5mm height (EMS Cat# 75959-03)
- Diatome Histo Diamond Knife (Diatome, Cat no. DH4540)
- Leica EM UC6 Ultramicrotome (Leica microsystems)
- Zeiss Gemini300 Field Emission Scanning Electron Microscopy with Gatan 3View (Carl Zeiss Microscopy GmbH) and a second generation FCC or relevant volume electron microscope

### Software

- DigitalMicrograph (Gatan)
- Fiji

### Reagent setup

- Complete DMEM: HG media containing 10% fetal bovine serum and 1X non essential amino acids.
- Complete DMEM: HG media containing 3% fetal bovine serum and 1X non essential amino acids.
- Fixation solution: 2.5% Glutaraldehyde, 2% paraformaldehyde in 0.1M Sodium Cacodylate Buffer (pH 7.2-7.4)
- TCH solution: 0.1g TCH (thiocarbohydrazide) to 10 mL ddH_2_O and place in a 60°C oven for 1 hour, agitate by swirling gently every 10 minutes to facilitate dissolving. Filter solution through a 0.22 μm filter right before use.
- 2% Osmium in ddH_2_O
- 1% Uranyl acetate in ddH_2_O
- Aspartic acid stock solution: dissolve 0.998g of L-aspartic acid in 250 mL ddH_2_O water. Note: aspartic acid dissolves more readily at pH 3.8. The solution is stable for 1-2 months at 4 °C.
- Walton’s lead aspartate solution^40^: dissolve 0.066g lead nitrate in 10 mL of the aspartic acid stock solution. Adjust the pH of the solution to 5.5 with 1M KOH.
- Durcupan ACM: Epoxy resin/ A (11.4 g); 964 hardener/ B (10 g); 964 accelerator/ C (0.3 g); Dibutyl phthalate / D (0.05-0.1 g)
- Hard Epon: EMbed 812 (5 mL); DDSA (2.25 mL); NMA (3 mL); BDMA (0.3 mL)

## PROCEDURE

### Culturing mammalian cells and sample fixation

<u>Step 1:</u> Culture adherent mammalian cells for 24 - 48 h in a 60 mm or 100 mm tissue culture dish for cell pellet, or 35 mm tissue culture dish with a gridded glass bottom for CLEM (final density 3-6 x 10^5^ cells) according to the standard protocol for the cell line. In our experiments, Vero cells were cultured in complete DMEM:HG media.

*Note:* For cell pellets, the cells can be collected when they reach 90% confluence. This ensures that there are enough cells to form a large cell pellet. For CLEM, the cells should be at 70% confluence or less as this will allow you to track the position of a cell of interest by both light and electron microscopy.

<u>Step 2:</u> For en face embedding of cells, proceed with light microscopy to identify specific cells of interest. Depending on the biological event being studied you can use brightfield or fluorescence microscopy to identify cells of interest. For cell pellet samples, proceed directly to step 4.

*Critical step:* 1) Image cells in an environmental chamber where temperature, humidity and CO2 concentrations are maintained 2) Capture images at high magnification (40X or 60X) and low magnification (5X or 10X) of multiple regions of interest using DIC and fluorescence imaging, as appropriate, to identify cells of interest. Images collected at high magnification will allow you to record both the cell shape and a more precise position relative to the surrounding cells. From images collected at low magnification, a map of the cell location showing the gridded lines, numbers or letters can be used as a marker when trimming the EM sample block, and searching the target cell of interest under EM. We recommend selecting 2 - 4 regions of interest per sample, that are located far apart from each other on the dish. This ensures that each region of interest will be reliably recovered during the subsequent processing steps when the sample is physically cut away from the polymerized resin.

<u>Step 3:</u> Discard tissue culture media and gently pipette 2 mL of fixative (2% PFA and 2.5% glutaraldehyde in 0.1M sodium cacodylate buffer (pH 7.2) to fix the cells. For cell lines that are more sensitive, add pre-warmed 1 mL 2 x fixative (4% PFA and 5% glutaraldehyde in 0.2M sodium cacodylate buffer, pH 7.2) directly into the dish with 1 mL of culture media.

*Critical step:* fixative solution should be freshly prepared

Note: the cells can be fixed first before light microscopy imaging. The fixative can be replaced with PBS and imaging can be done under the light microscope without an environmental chamber.

*Caution:*Please refer to the material safety data sheet of each compound to ensure its careful handling.

<u>Step 4:</u> For cell pellet fixation, after 1 min of fixation with 2% paraformaldehyde and 2.5% glutaraldehyde in 0.1M sodium cacodylate buffer (pH 7.2), detach cells from cell culture dish using cell scraper. Transfer the cells into a 1.5 mL tube and centrifuge immediately at 5000 rpm for 2 min. Rotate the tube 180 degrees and centrifuge at 5000 rpm for an additional 2 min. Cells can be observed as a pellet at the bottom of the tube.

Critical step: 1) scrape cells in a single direction, such that you observe a sheet of cells coming off the dish. 2) in-dish fixation should be limited to less than 3 minutes to avoid disintegration of the cell monolayer which can make recovery of the dense cell pellet during centrifugation tough.

Step 5: Change to 1 mL fresh fixative (2% paraformaldehyde and 2.5% glutaraldehyde in 0.1M sodium cacodylate buffer (pH 7.2) and continue to fix the cells overnight at 4 °C.

### Sample staining and embedding

<u>Step 6:</u> Discard fixative and add 1 mL 0.1M cacodylate buffer for 10 min at room temperature. Repeat this buffer rinsing step two more times.

Step 7: Discard remaining buffer and incubate the sample in a freshly made 200 uL reduced osmium solution that contains 2% osmium, 1.5% potassium ferrocyanide in 0.1M sodium cacodylate buffer, for 1.5 h at RT in dark.

*Caution:* do not disturb the cell pellet during all sample processing steps. Toxic/ carcinogenic/ Please refer to the material safety data sheet of the compound to ensure its careful handling.

<u>Step 8:</u> Wash the sample in 1 mL ddH_2_O for 3 min at room temperature. Repeat this wash step four more times

Step 9: Incubate the sample in freshly prepared 1 mL TCH solution (1%) for 20 min at room temperature.

<u>Step 10:</u> Wash the sample in 1 mL ddH_2_O for 3 min at room temperature. Repeat this wash step four more times

Step 11: Incubate the sample in 200 uL 2% Osmium in ddH_2_O for 40 min at room temperature.

<u>Step 12:</u> Wash the sample in 1 mL ddH_2_O for 3 min at room temperature. Repeat this wash step four more times

Step 13: Incubate the sample in 0.5 mL 1% uranyl acetate in aqueous solution, overnight at 4 °C in the dark

*Caution:* Toxic/ carcinogenic Please refer to the material safety data sheet of the compound to ensure its careful handling.

<u>Step 14:</u> Wash the sample in 1 mL ddH_2_O for 3 min at room temperature. Repeat this wash step four more times

Step 15: Incubate samples in 1mL of *En bloc* Walton’s lead aspartate staining for 30 min inside a 60 °C oven.

*Critical step:* Walton’s lead aspartate solution should be prewarmed at 60 °C for 30 min. Avoid precipitate formation.

*Caution:* Please refer to the material safety data sheet of each compound to ensure its careful handling.

<u>Step 16:</u> Wash the sample in 1 mL ddH_2_O for 3 min at room temperature. Repeat this wash step four more times

<u>Step 17:</u> Carry out sample dehydration in a graded ethanol series as follows (1 mL per tube, or 2 mL per 35 mm dish)

30% ethanol in ddH_2_O; 1X for 10 min at 4 °C

50% ethanol in ddH_2_O; 1X for 10 min at 4 °C

70% ethanol in ddH_2_O; 1X for 10 min at 4 °C

85% ethanol in ddH_2_O; 1X for 10 min at 4 °C

95% ethanol in ddH_2_O; 1X for 10 min at 4 °C

100% ethanol; 1X for 10 min at 4 °C

100% ethanol; 2X for 10 min at room temperature

<u>Step 18:</u> Carry out resin infiltration of the sample using a graded Durcupan resin series as follows (0.5 mL per tube, or 1 mL per 35 mm dish)

25% Durcupan in ethanol; 1X for 4 h at room temperature

50% Durcupan in ethanol; 1X for overnight at room temperature

75% Durcupan in ethanol; 1X for 2 h at room temperature

100% Durcupan; 1X for 2 h at room temperature

100% Durcupan; 2X for 1 h at room temperature

*Note:* For cell pellet embedding, a hard formulation of EMbed^41^ can replace Durcupan, which is easier for sectioning with an ultramicrotome. However, EMbed will dissolve the cell culture dish, therefore it is not appropriate for an *en face* CLEM project.

<u>Step 19:</u> For sample embedding of *en face* samples, tilt the dish at a 45° angle in order to discard as much leftover resin from the dish as possible. Add a thin layer of 100% Durcupan resin (∼100 ul) directly onto the glass coverslip. For cell pellets, there are two options when sample embedding, 1) use a wood applicator stick to transfer the sample to an embedding mold, or 2) directly embed the sample inside the tube if the cell pellet is too small. With either sample embedding method, add 100% Durcupan to fill the embedding mold or ∼200 uL to the tube if directly embedding the sample inside the tube. Place embedded samples in a 60 °C oven for 48 - 60 h to allow resin-infiltrated samples to polymerize and harden.

*Critical step:* 1) for *en face* samples, be sure to add a requisite volume of 100% Durcupan (one or two drops) that will cover just the glass coverslip and not the surrounding areas of the dish and 2) for cell pellet samples in which the sample is very small, sample embedding directly in the tube is recommended.

*Note:* Annotate samples clearly to prevent any confusion downstream when working with multiple samples. For *en face* samples, write down a sample number on a piece of paper and place it on the plastic area of the dish, so that any leftover resin will stick the number tightly to the dish. For cell pellet samples, put a sample number into an embedding mold without it overlapping with the sample, or stick it to the inner wall of the tube.

### Sample trimming and mounting

For cell pellet samples:

Step 20: Bend the sample embedding mold, or cut open a tube using a razor blade to take out a polymerized sample block.

<u>Step 21:</u> Lock the sample block on the specimen holder of the microtome, then trim the sample block under the microtome. Trim one edge of the sample block containing the cell pellet using a razor blade to expose the cell sample.

<u>Step 22:</u> Place the sample block in the specimen arc of the ultramicrotome and section the exposed sample surface using a Diatome Histo Diamond knife to create the blockface. The knife holder and specimen arc angles should be set to zero.

<u>Step 23:</u> Take the sample block out of the specimen arc. Using a razor blade, trim the remaining edges of the sample block into a shaped pillar.

*Critical step:* Whilst trimming the sample block edges, ensure that: 1) the excess resin is trimmed away on the remaining edges in order to expose the sample and 2) each side of the pillar is of an equal dimension relative to the block face.

Step 24: Place the sample block under a stereomicroscope and cut away the newly created 3 view sample blockface pillar with a razor blade.

*Note:* Laying the block flat and protecting the pillar with scotch tape before cutting, will help to ensure that the pillar is not lost. Doing so on a piece of white paper will contrast the osmium-darkened sample, making the pillar easier to see.

Step 25: Glue the pillar to a 3View SEM pin stub (EMS Cat# 75959-02) using silver conductive epoxy.

*Critical step:* Whilst gluing the sample, ensure that 1) the sample block face created using a Diatome Histo Diamond knife is glued directly to the surface of the 3View SEM pin stub, with the side cut away from the sample block facing up and 2) each side of the pillar is also covered by silver epoxy.

Step 26: Place the 3View pin stub in a 60 °C oven overnight to polymerize the epoxy.

Step 27: Place the pin stub in the specimen arc of the ultramicrotome and section the exposed sample surface using a Diatome Histo Diamond knife, where both the knife holder and specimen arc are set to angles of zero.

Step 28: Remove the pin stub from the ultramicrotome and trim away any excess epoxy from the sides of the pillar.

For *en face* samples:

<u>Step 29:</u> Freeze the dish with the resin embedded sample in liquid nitrogen to carefully separate the glass coverslip from the polymerized cell monolayer. Repeat these freeze thaw cycles if necessary.

*Critical step:* Immediately after freezing, use a very sharp razor blade (EMS Cat# 72000-WA) to separate the perimeter of the circular coverslip from the dish, by lodging the razor’s edge between the perimeter of the coverslip and dish while rotating the dish.

Step 30: Identify the region of interest (ROI) and its respective, letter-number location square, within the location-based grid, on the sample side of the polymerized monolayer (Figure 3A).

Step 31: On the sample side mark off a square around the letter-number location containing the ROI, using a razor blade (Figure 3B).

*Critical step:* Take care not to damage the surface of the monolayer-ROI during this and all subsequent steps, as it is no longer protected by the glass coverslip; simply touching it may destroy the sample and/or its respective location-specific letter/number.

Step 32: Turn over the polymerized sample monolayer and mark the location of the ROI with an “x” on the non-sample side of the resin; use the square marked by a razor on the sample side as a reference in doing this (Figure 3B).

Step 33: Using a stereomicroscope, confirm that the “x” on the non-sample side, aligns with the ROI on the sample side (Figure 3B).

<u>Step 34:</u> Carefully place a 1 x 2 mm slot grid over the ROI on the sample side. Ensure that the ROI is centered within the slot. Using a razor blade, place marks at the sides of the slot grid so that the marks are parallel to the 2 mm side of the slot. These marks, in combination with the marks made in step # 3, should create a rectangle equal to the diameter of the slot grid (Figure 3B).

Step 35: Using a razor separate this rectangle, one side at a time, from the larger polymerized monolayer (Figure 3B).

*Critical step:* Confirm that the non-sample side of the rectangular resin piece contains the centered “x” from step 30 (Figure 3B).

Step 36: Set the rectangular resin piece aside, either in a microcentrifuge tube or a small gelatin capsule, so as not to lose the sample during subsequent steps.

Step 37: Using a conductive silver epoxy, glue two slot grids one on top of the other to the top of a 3View SEM pin stub (EMS Cat# 75959-03) (Figure 3B).

*Note:* Make sure that the slots of both slot grids are aligned

Step 38: Place the 3View SEM/ slot grid pin stub into a 60°C oven overnight to polymerize the epoxy.

*Note:* Once dried, remove the now polymerized, excess silver epoxy from the interior of the superimposed slots using the edge of a razor or scalpel.

Step 39: Glue the ends of the rectangular resin piece to the broader parts of the slot grid, sample side down, using the same conductive epoxy.

*Critical step:* Ensure that the ROI target is centered in the slot of the slot grid and that the “X” marked on the non-sample side in Step 30 faces up. Be very careful to avoid getting epoxy on the ROI itself by placing a piece of tape on the sample side.

<u>Step 40:</u> Allow the glue between the 3View SEM/ slot grid pin stub and the rectangular resin piece to fully polymerize. Leave samples in a 60 °C oven overnight.

Step 41: Place the 3View SEM/ slot grid pin stub into the specimen arc of the ultramicrotome, taking care to ensure that both the specimen arc and knife holder are at zero degrees.

<u>Step 42:</u> Using a Diatome Histo Diamond knife, section the now exposed non-sample side, at 500 nm section thickness and speed 1mm/second. Continue to section until most of the resin from the non-sample side is removed and the rectangle is as thin as possible.

*Critical step:* Thinning the non-sample side of the rectangular resin piece not only removes excess resin in order to increase the conductivity of the sample, but also ensures that the sample is parallel to the 3View SEM pin stub surface.

Step 43: Using a sharp razor, very carefully remove the thinned resin rectangle from the 3View SEM/ slot grid pin stub.

Step 44: Under a stereomicroscope, cut the ends of the rectangle away so that the resin piece becomes a square with the ROI centered.

Step 45: Glue this final square resin piece to the 3View SEM pin stub (EMS Cat# 75959-02), sample side up, using a conductive silver epoxy (Figure 3B).

Step 46: Place the 3View pin stub in a 60 °C oven overnight, to polymerize the epoxy.

*Critical step:* Sample on 3View pin can be baked in a 100 °C oven for 1 h before sputter coating to increase the stability of the sample during SBF cutting.

<u>Step 47:</u> Sample pin is loaded into a sputter coater. A layer of gold 15 nm thickness is sputtered onto the sample block faces. This improves conductivity and the reflective surface aids in the manual sample approach detailed in step 53.

### Sample loading

<u>Step 48:</u> Sample pin is slotted and secured into the 3View microtome chuck by tightening a set screw. The sample is first rotated inside of the chuck such that the leading face of the block is rotated 90 degrees clockwise from the set screw. This is done with the assistance of a stereo microscope.

<u>Step 49:</u> Check that the sample is loaded and secured into the microtome. The diamond knife is shifted laterally and the cutting window position adjusted to center over the span of the block face. Using a stereo microscope and the reflection of the knife edge on the block face, manually approach the sample to the cutting plane.

*Critical step:* Care should be taken to avoid contact between the block face and the knife edge as this will leave an imprint several microns deep. A practiced user should get the block face 5-10 µm away from the cutting plane.

*Note:* Leave the sample inside SEM chamber overnight before imaging to allow for a more thorough off-gassing and temperature equalization of the hardware and sample

### Sample imaging

Step 50: Adjust the stage position using the DigitalMicrograph software (Gatan).

<u>Step 51:</u> Perform coarse cuts at 100 nm slice thickness to approach the sample blockface to the ultramicrotome knife. Acquire low magnification reference scans at intervals to confirm that the sample block is in the cutting plane of the knife.

<u>Step 52:</u> For cell pellet embedded samples, acquire a low resolution scan that captures the sample blockface. Proceed directly to step 55.

<u>Step 53:</u> For *en face* embedded samples, acquire an image with the secondary electron detector. This provides a clearer view of the sample blockface surface and makes visible the debossed grid numbers and borders.

Step 54: Use low magnification images from light microscopy and the secondary electron detector to correlate the position of the cell of interest on the block face.

Step 55: Create and position an imaging tile (or tile-set) over the cell of interest.

<u>Step 56:</u> Set the chamber pressure, aperture size and electron beam settings. Refer to parameters listed in table1.

<u>Step 57:</u> Using the secondary electron detector, activate Beam Wobble in SmartSEM (Carl Zeiss GmbH). Adjust the aperture X and Y position to center and align the mid-lens aperture. Once the image is sufficiently still the aperture is aligned and Beam Wobble can be deactivated.

<u>Step 58:</u> In Gatan Digital Micrograph, edit the image tags in the Global Info settings to assign any metadata that is not automatically generated. These tags include sample name, lab or principle investigator name, sample block number (if applicable), embedding resin used, aperture size and FCC pressure. Imaging parameters such as slice thickness, pixel size, dwell time and acceleration voltage are automatically embedded in the metadata.

<u>Step 59:</u> Assign a storage directory for the saved dataset. Each scan is saved as an individual file in the native Gatan Digital Micrograph ‘.dm3’ file format. Multiple regions of interest are saved into their respective folders (ROI_00, ROI_01, et cetera).

<u>Step 60:</u> Begin the dataset collection. For multiple regions of interest, the software will ask to confirm the positioning of the scans. The software will also ask the user to assign a region as a “monitor stack”, which allows for the storage of several sequential scans in RAM for quick viewing. This is used to monitor cutting quality, sample charge rate and to perform any focus corrections during data collection.

Step 61: Adjust focus and stigmation as the first image is being acquired.

*Critical step:* It is advised to monitor the first scans to assess whether there are any sample charge artifacts in the image. FCC pressure can be increased to compensate for sample charge or reduced to improve image contrast if the sample is sufficiently conductive.

Step 62: Monitor imaging periodically to adjust any focus drift that may occur during acquisition.

## TROUBLESHOOTING

Troubleshooting advice can be found in table 3.

**Table 3:**
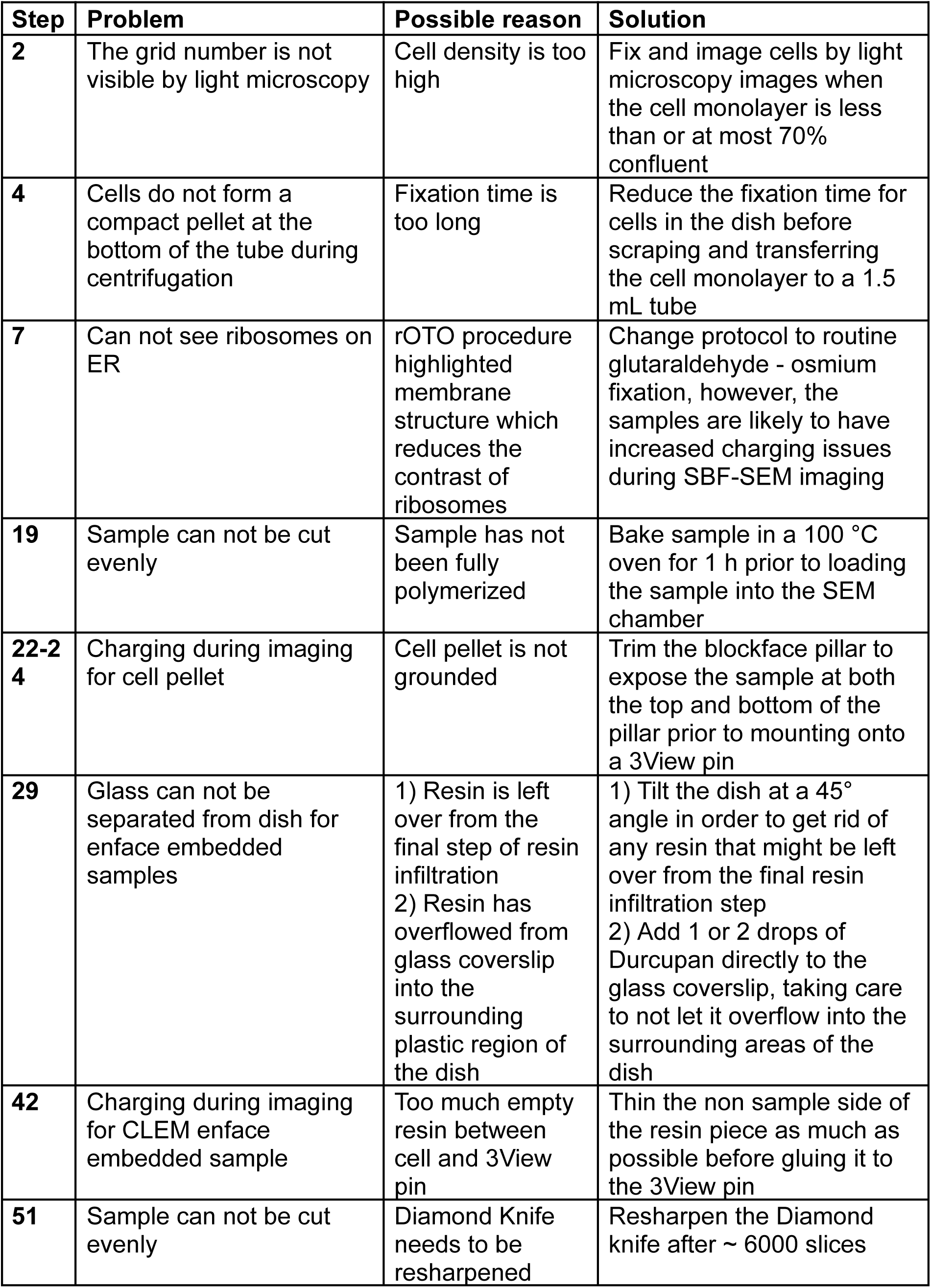
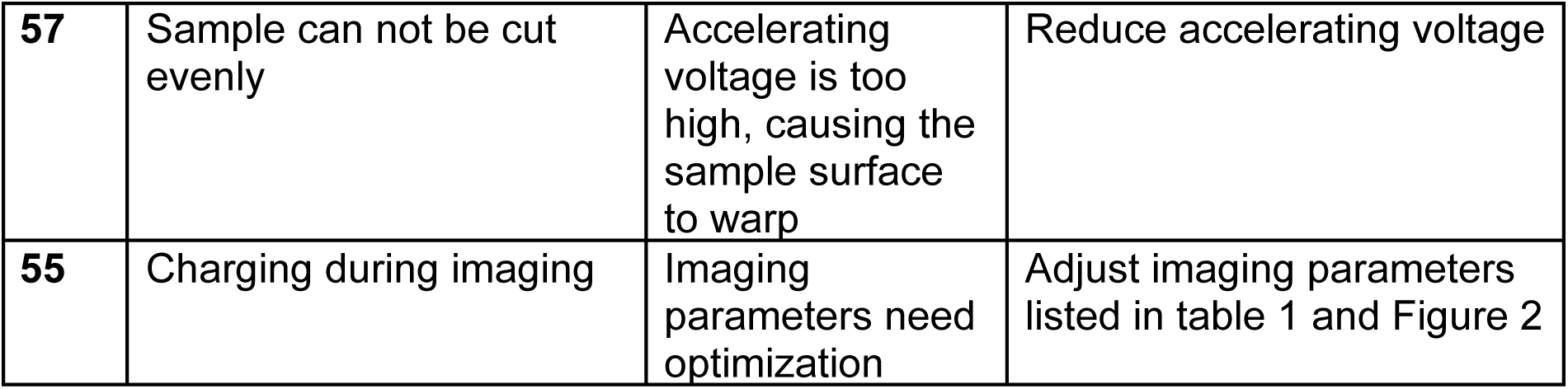
Troubleshooting table.

| **Step** | **Problem** | **Possible reason** | **Solution** |
| --- | --- | --- | --- |
| **2** | The grid number is not visible by light microscopy | Cell density is too high | Fix and image cells by light microscopy images when the cell monolayer is less than or at most 70% confluent |
| **4** | Cells do not form a compact pellet at the bottom of the tube during centrifugation | Fixation time is too long | Reduce the fixation time for cells in the dish before scraping and transferring the cell monolayer to a 1.5 mL tube |
| **7** | Can not see ribosomes on ER | rOTO procedure highlighted membrane structure which reduces the contrast of ribosomes | Change protocol to routine glutaraldehyde - osmium fixation, however, the samples are likely to have increased charging issues during SBF-SEM imaging |
| **19** | Sample can not be cut evenly | Sample has not been fully polymerized | Bake sample in a 100 °C oven for 1 h prior to loading the sample into the SEM chamber |
| **22-24** | Charging during imaging for cell pellet | Cell pellet is not grounded | Trim the blockface pillar to expose the sample at both the top and bottom of the pillar prior to mounting onto a 3View pin |
| **29** | Glass can not be separated from dish for enface embedded samples | 1) Resin is left over from the final step of resin infiltration<br>2) Resin has overflowed from glass coverslip into the surrounding plastic region of the dish | 1) Tilt the dish at a 45° angle in order to get rid of any resin that might be left over from the final resin infiltration step<br>2) Add 1 or 2 drops of Durcupan directly to the glass coverslip, taking care to not let it overflow into the surrounding areas of the dish |
| **42** | Charging during imaging for CLEM enface embedded sample | Too much empty resin between cell and 3View pin | Thin the non sample side of the resin piece as much as possible before gluing it to the 3View pin |
| **51** | Sample can not be cut evenly | Diamond Knife needs to be resharpened | Resharpen the Diamond knife after ~ 6000 slices |
| **57** | Sample can not be cut evenly | Accelerating voltage is too high, causing the sample surface to warp | Reduce accelerating voltage |
| **55** | Charging during imaging | Imaging parameters need optimization | Adjust imaging parameters listed in table 1 and Figure 2 |

## TIMING

This protocol can be performed over multiple days. Once samples are embedded in resin they are stable at room temperature indefinitely once infiltrated by and polymerized in resin.

<u>Step 1:</u> Culturing mammalian cells, plus an additional 24 - 48 h for cells to grow

<u>Step 2:</u> Brightfield or fluorescence microscopy for CLEM workflow with *en face* samples, 1 - 2 h

Steps 3-5: 2 h fixation at room temperature plus an additional 24 h at 4 °C.

Step 6-19: Sample staining, embedding and polymerization, 5 days plus any additional time for reagent setup and clean up.

Step 20-28: Cell pellet sample trimming and mounting, 2 h plus any additional time for epoxy polymerization, reagent setup and clean up.

Steps 29-47: *en face* sample trimming and mounting, 4 h plus any additional time for epoxy polymerization, reagent setup and clean up.

Steps 48-49: sample loading on the SBF-SEM, 1 h plus an additional 12 h to let the system equilibrate.

Steps 50-62: sample imaging, several hours to days depending on the number of ROIs being collected, the volume assigned as well as pixel size and dwell time.

## ANTICIPATED RESULTS

### Representative results from imaging tissue culture cells by SBF-SEM

Using the protocol described above, a new user should be able to successfully fix, stain, embed, trim, mount and image tissue culture cells using either a cell pellet or an *en face* protocol. In this section we show representative results from imaging a cell pellet embedded sample and an *en face* embedded sample (Figure 5). We have used these protocols to investigate the biology of an obligate intracellular parasite, *E. intestinalis* in Vero cells^42^. In both the cell pellet and *en face* examples described here, the goal is to acquire high resolution images in which the ultrastructures of organelles like the mitochondria and/ or endoplasmic reticulum can be resolved. We used a pixel size of 2 nm. However, a 4-6 nm pixel size would be a reasonable starting point in order to answer a majority of biological questions that require a well defined organelle structure. The resulting image stacks can be examined using freely available image analysis software, such as Fiji^38^, and more complex analyses such as segmentation of cellular structures and 3D reconstruction can be carried out using dedicated software packages, such as Dragonfly^39^.

**Figure 5:**
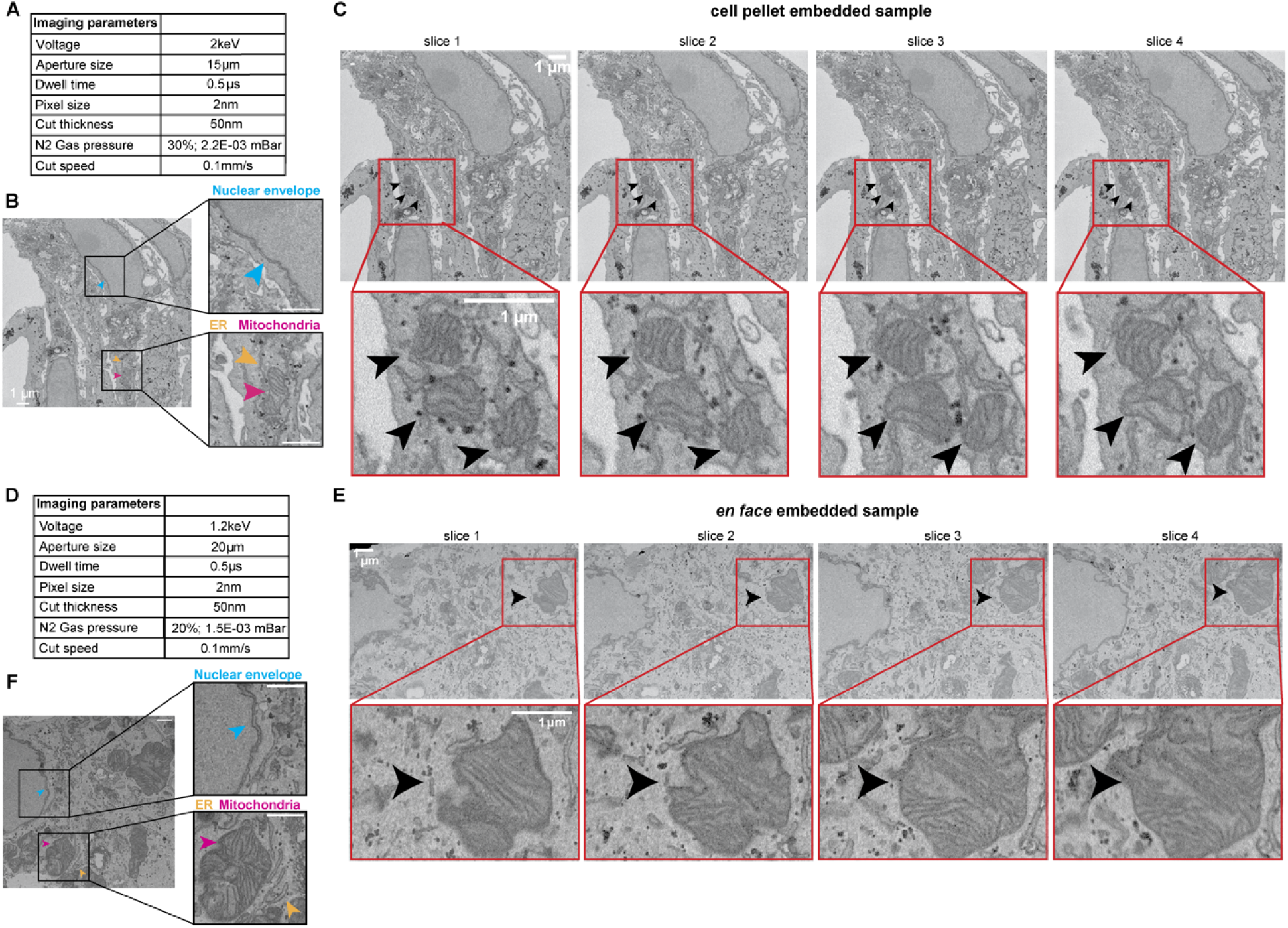
Representative imaging results for cell pellet and *en face* embedded samples. (A, D) Tables showing the imaging parameters used to achieve good signal-to-noise and avoid charging artifacts in the cell pellet sample (A) or *en face* sample (D). (B, E) Representative slices from the cell pellet (B) or en face (E) samples using the imaging parameters listed in (A) and (D), respectively. Insets highlight the ultrastructure of different host cell organelles - nucleus, blue arrowhead; mitochondria, magenta arrowhead; ER, yellow arrowhead. (C, F) Slices 1-4 of the cell pellet (C) or en face (F) samples. Scale bars 1 μm.

## Supporting information

Movie 1: Image stack of the cell pellet embedded sample imaged in Figure 5.

Movie 2: Image stack of the en face embedded sample imaged in Figure 5.

## Acknowledgements

We thank Jason (Xiangxi) Liang from the NYU Microscopy Core with assistance in the preparation of EM samples. We thank Ari Davydov for critical reading and feedback on the manuscript and all members of the Bhabha / Ekiert labs for helpful discussions. We gratefully acknowledge the following funding sources: SSP-2018-2737 (Searle Scholars Program, to G.B.), R01AI147131 (NIAID, to G.B.), Irma T. Hirschl Career Scientist Award (to G.B.). G.B. is a Pew Scholar in the Biomedical Sciences, supported by The Pew Charitable Trusts (PEW-00033055). The NYU Microscopy Core is partially supported by NYU Cancer Center Support Grant NIH/NCI P30CA016087, and Zeiss Gemini 300 SEM with 3View was purchased with support of NIH S10 ODO019974.

## Movie Legends

**Movie 1:** Image stack of the cell pellet embedded sample imaged in Figure 5.

**Movie 2:** Image stack of the *en face* embedded sample imaged in Figure 5.

